# Heart Murmur Detection in Phonocardiogram Data Leveraging Data Augmentation and Artificial Intelligence

**DOI:** 10.1101/2025.07.16.665251

**Authors:** Melissa Valaee, Shahram Shirani

## Abstract

**Background:** With a 17.9 million annual mortality rate, cardiovascular disease is the leading global cause of death. As such, early detection and disease diagnosis are critical for effective treatment and symptom management. Cardiac auscultation, the process of listening to the heartbeat, often provides the first indication of underlying cardiac conditions. This practice allows for the identification of heart murmurs caused by turbulent blood flow. In this exploratory research paper, we propose an AI model to streamline this process that improves diagnostic accuracy and efficiency.

**Methods:** We utilized data from the 2022 George Moody PhysioNet Heart Sound Classification Challenge, comprised of phonocardiogram recordings of individuals under 21 years of age in Northeast Brazil. Only patients who had recordings from all four heart valves were included in our dataset. Audio files were synchronized across all recordings and converted to Mel spectrograms before being passed into a pre-trained Vision Transformer, and finally a MiniROCKET model. Additionally, data augmentation was conducted on audio files and spectrograms to generate new data, bringing our total sample size from 928 spectrograms to 14,848.

**Results:** Compared to existing methodologies, our model yielded significantly enhanced quality assessment metrics, including Weighted Accuracy, Sensitivity, and F-Score, as well as a faster evaluation speed of 0.02 seconds per patient.

**Conclusion:** The implementation of our method for the detection of heart murmurs can supplement physician diagnosis and contribute to earlier detection of underlying cardiovascular conditions, faster diagnosis times, increased scalability, and enhanced adaptive abilities.

## INTRODUCTION

At an estimated 17.9 million, cardiovascular diseases are the leading cause of death globally^1^. Early detection of such diseases is critical in their treatment and management. One critical diagnostic test for the detection of underlying cardiac conditions is cardiac auscultation. This practice involves listening to the heartbeat to identify heart murmurs caused by turbulent blood flow, which may be indicative of underlying complications^2^. Auscultation is conducted in routine physical examination and is traditionally carried out with a stethoscope^2,3^. Unfortunately, this method has limited sensitivity and accuracy due to high inter-rater discrepancy, even among trained healthcare professionals^3^. Given the importance of heart murmur detection in the diagnostic process, an enhanced protocol is necessary.

In recent years, evidence-based research regarding the applications of artificial intelligence in medical practices has demonstrated increased promise as a way to streamline and supplement physician diagnosis. Artificial intelligence has been implemented in various medical settings to enhance patient care through improved diagnostic capabilities, streamlined workflow and reduced medical errors and costs^4^. However, artificial intelligence is also limited in that its efficacy is heavily reliant on the data used to train the model. To ensure efficacy, algorithms must be trained on vast quantities of high-quality data. Outdated, imbalanced, or biased data may result in severe compromises^5^.

To address the aforementioned limitations in cardiac auscultation, we propose an artificial intelligence model that is able to detect heart murmurs with high accuracy and speed. Our model, leveraging data augmentation and machine learning, demonstrates increased Weighted Accuracy, Sensitivity, Specificity, Negative Predictive Value, Positive Predictive Value, and F-Score compared to pre-existing methods, while allowing for faster processing and classification times.

## METHODS

Our heart murmur detection method, as outlined in Figure 1, uses Multivariate Minimally Random Convolutional Kernel Transform (MiniROCKET) as its foundation^6^. First, phonocardiogram data from the 2022 PhysioNet Heart Sound Classification Challenge Database was filtered to only include trials that had recordings of all four valves^7,8^. Each set of four phonocardiogram recordings underwent a series of 10 random time-series data augmentation techniques applied a total of four times. Subsequently, a Mel spectrogram was generated from each audio file. For each of these spectrograms, one random data augmentation technique is applied three times to generate a total of three augmented spectrograms from each (Figure 2). All spectrograms are then flattened and passed into a pre-trained Vision Transformer and a vector of logits is returned. Then, a set of fixed kernels is convolutionally passed over the logit vector, finding the dot product of the kernel and the data. The product is then represented as a feature map that is input to a ridge regression model that completes the classification. Our classifier yields three results: “murmur is present”, “murmur is absent”, or “further assessment required” – if it can not make the classification with sufficient certainty.

**Figure 1.**
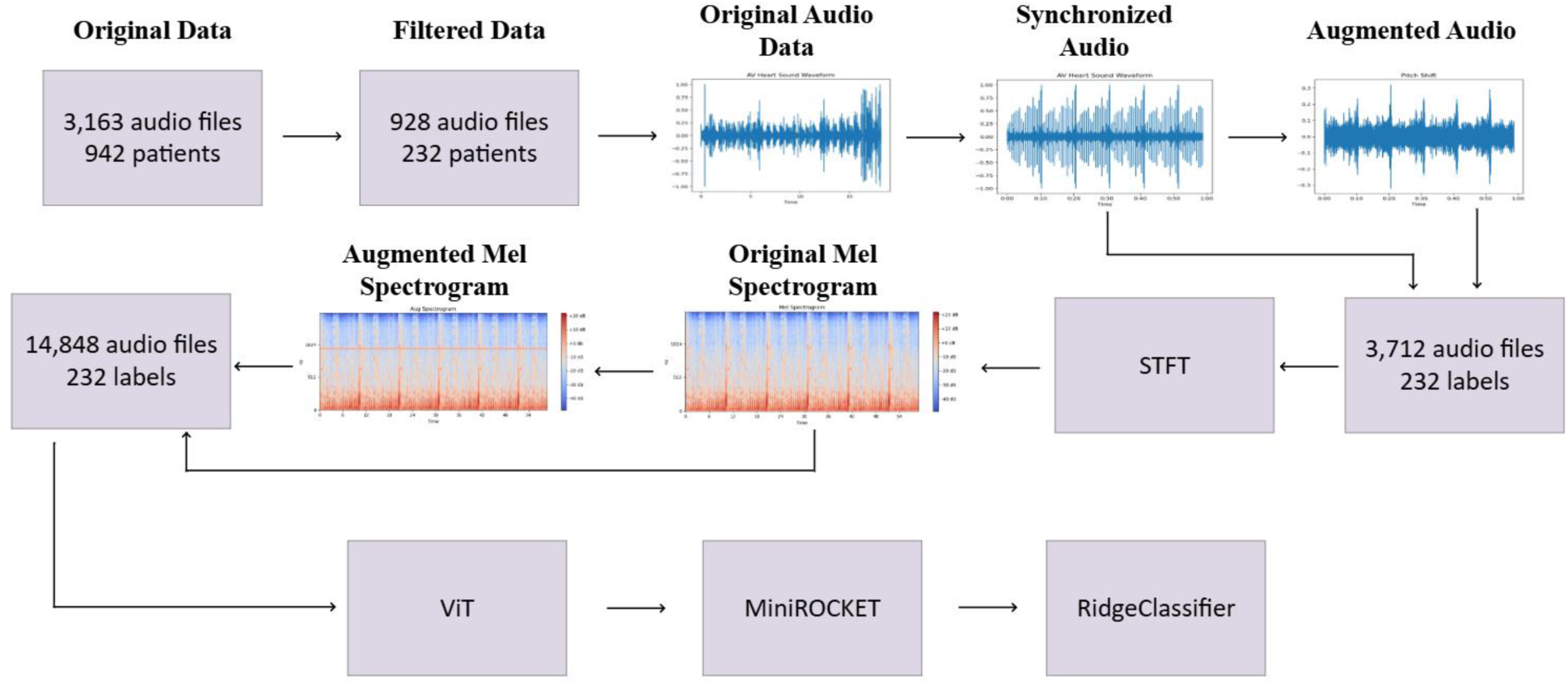
A visualization of our methodology, beginning with original sound data, which was then filtered such that only patients with data from all four heart valves were used. The heartbeats are then temporally aligned, and various augmentation techniques are implemented to generate more audio data. Then, a Mel spectrogram is generated from all data (real and augmented). Subsequently, a number of augmented spectrograms are generated from the original spectrogram. Each Mel spectrogram is then passed through the Vision Transformer, which generates the logit vector that is passed to the MiniROCKET model. Finally, a Ridge Classifier is used for classification.

**Figure 2.**
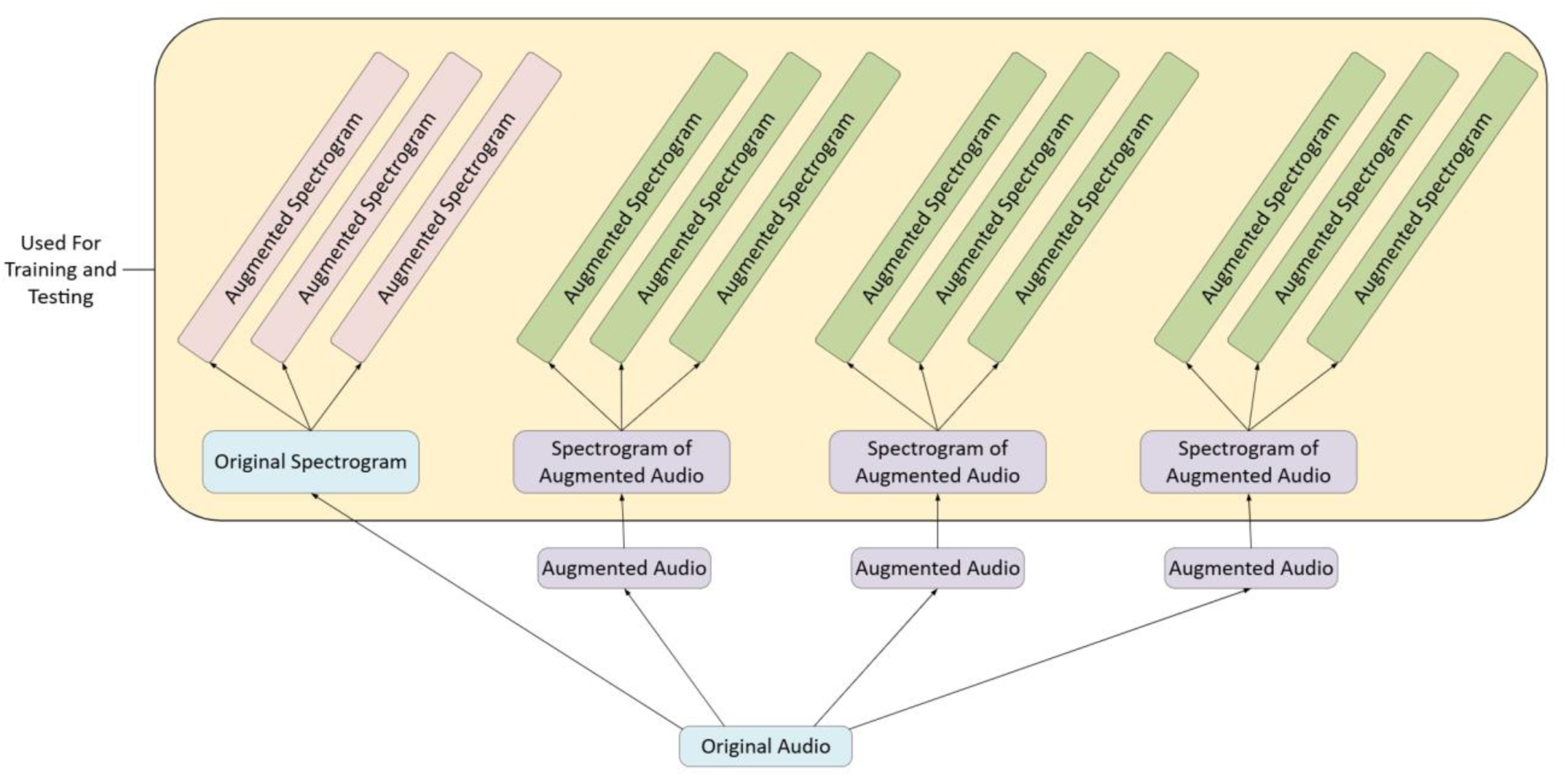
A visualization of the Data Augmentation process. Each original PCG audio file yielded three augmented audio files. From each audio file, one Mel spectrogram was generated, which served as the template for three additional augmented spectrograms.

## DATA AVAILABILITY

All data used throughout the training and testing phases of our model development was obtained from the CirCor DigiScope Phonocardiogram Dataset for the George B. Moody PhysioNet Challenge 2022 on Heart Murmur Detection from Phonocardiogram Recordings^7^. This publicly available dataset contains 5272 sound recordings from 1568 anonymized patients aged 0 to 21 years collected in Northeast Brazil from July to August of 2014 and June to July of 2015. The dataset contains recordings from all primary auscultation points: the Aortic Valve, the Pulmonic Valve, the Tricuspid Valve, and the Mitral Valve; however, not all patients have recordings for all four valves. Furthermore, each audio file was labelled based on whether or not a murmur was present at a given location. We included only patients who had data from all four valves, grouping them together and assigning a label based on whether a murmur is present at any of the four valves. Thus, we began with a sample size of 928 audio files spanning 232 labels (each labeled patient consists of four audio recordings, one for each valve.

## AUDIO SYNCHRONIZATION

In our methodology, we leverage Multivariate MiniROCKET, a supervised machine learning algorithm designed to rapidly and accurately classify data based on multiple directly correlated input channels^6^. For our implementation, audio from each valve is considered its own channel, and they must be made congruous through heartbeat synchronization. At a given time, two audio signals are compared to each other; the heartbeat peaks are identified and lined up such that the first heartbeat of each audio file occurs at the same timestamp. This process is carried out until all four audio signals are aligned with each other. Then, each individual signal is looped until it reaches a time of 58 seconds – the length of our longest audio file – to ensure that all model inputs are the same length.

## DATA AUGMENTATION

### Audio File Augmentation

Data augmentation is a process that generates new samples from existing data^9^. This task is accomplished by applying various transformations to the original dataset, such as rotation, flipping, and noise injection. In applying these transformations, the model is able to learn from a more diverse set of inputs, thus increasing its robustness and generalizability.

Data augmentation is particularly beneficial in situations where collecting labelled data is challenging, expensive, or time-consuming^10^. Through data augmentation, the dataset can be expanded, which allows the model to more easily and accurately identify patterns in data, and thus, yields improved accuracy when exposed to new data. Additionally, data augmentation promotes data diversity; thus the model may learn to better handle variations in data^11^.

Furthermore, data augmentation helps prevent overfitting of the model. Overfitting occurs when a model simply memorizes the training data rather than identifying the overarching patterns. In these instances, the model performs poorly on unseen data. However, introducing variations in the training data through data augmentation forces the model to learn generalized features, thus, reducing the risk of overfitting^9,11^.

We implemented various data augmentation techniques at two distinct stages in our methodology. We based our augmentation techniques on those commonly used in existing literature^9^. Firstly, we applied ten random transformations from the following list to each audio file: Gaussian noise injection, pitch shift, time shift, and time warp (Figure 3).

**Figure 3.**
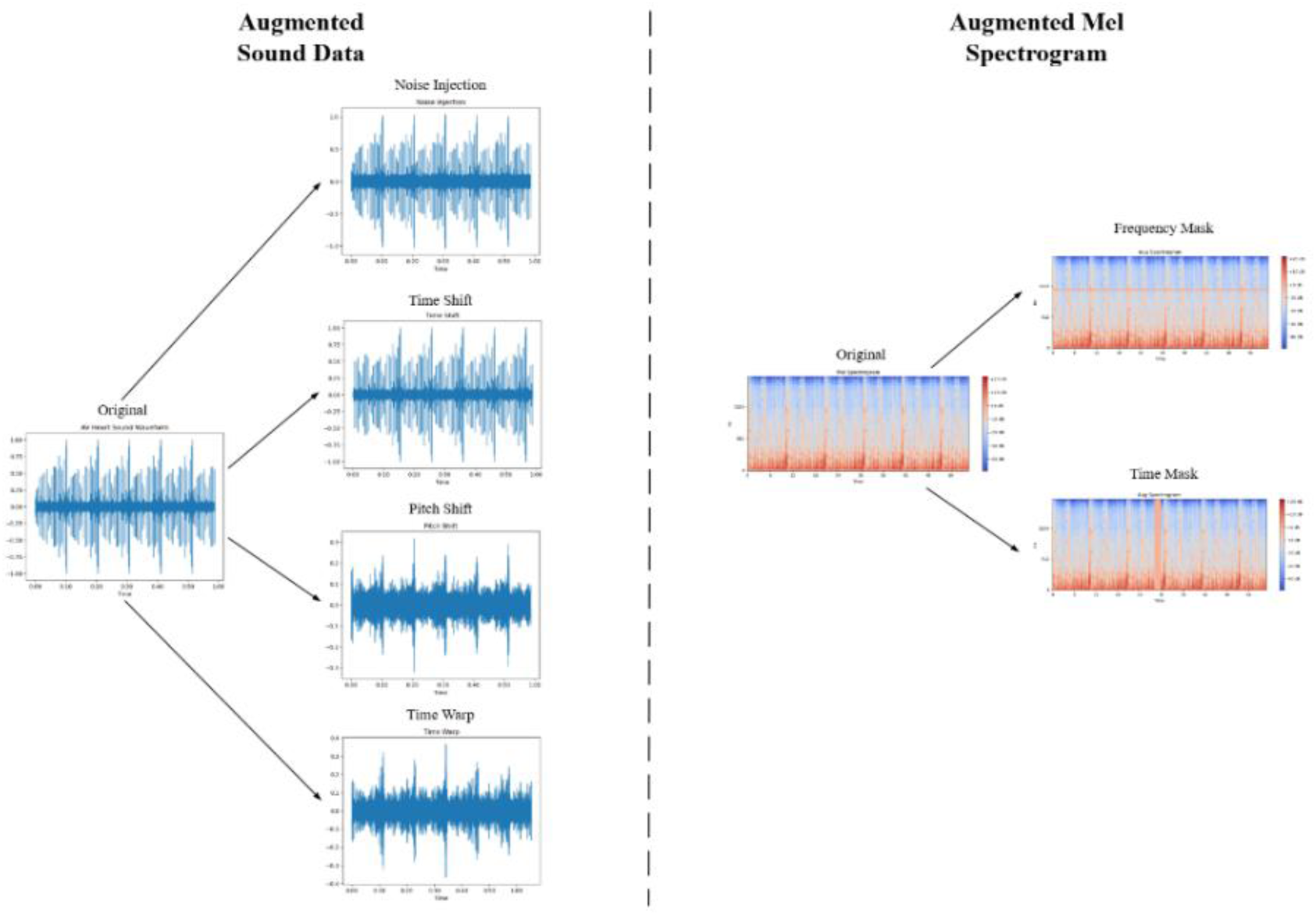
A visualization of the implemented data augmentation techniques. For sound data (left), we implemented noise injection, time shift, pitch shift, and time warp. For Mel spectrograms (right), we implemented frequency masking and time masking.

#### Gaussian Noise Injection

Gaussian noise is a type of statistical noise that follows a bell-shaped normal distribution. It is common in machine learning applications^9^. First, a sequence of random numbers is generated such that the length of this sequence is equal to the length of the input data, the mean of the sequence is 0, and its standard deviation is 1. It is then scaled by a noise factor and added to the input data in an element-wise fashion. Thus, each element in the noise sequence is summed with the corresponding index of the data signal sequence.

#### Pitch Shift

Pitch refers to the perceived frequency of a sound, with higher pitch sounds corresponding to higher frequency. First, the spectrogram containing the time-frequency representation of the audio file is generated. Then, pitch shifting is completed by manipulating the frequencies according to the following formula

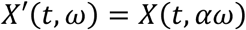

where *X(t, ω)* is the original frequency representation, *α* is the pitch-shifting factor (*α* > 1 increases the pitch), and *X’(t, ω)* is the pitch-shifted frequency representation. During this process, all frequency components are shifted together (both the fundamental frequency and all associated harmonics), thus preserving the sound of the original data. Finally, the audio signal is reconstructed from the augmented spectrogram.

#### Time Shift

Time shift causes a randomly generated unidirectional shift in the data. This is achieved by shifting the values in each index of the data signal. The order of the data is preserved, but each point is moved to a different timestamp. In our implementation, the values at the ends of the signal are wrapped around at the boundary to preserve the signal upon shifting.

#### Time Warping

Time warping alters the temporal characteristics of the audio file while preserving the original pitch. To achieve this, the audio file is first converted to the time-frequency by a Short Time Fourier Transform (STFT). An STFT converts the audio signal into a spectrogram, which is a visual representation of the audio signal’s frequency over time. In this representation, the time domain can be manipulated independently of pitch. Finally, the audio file is restored to its initial representation through an inverse STFT.

All transformations were applied across all four data channels to preserve their correlation. From each patient’s original data, we generated three sets of augmented audio data, bringing our total number of sound recordings to 3,712 audio files from our original 928 files.

### Mel Spectrogram Augmentation

Subsequently, a Mel spectrogram is utilized to more closely mimic human auditory perception. To achieve this representation, each audio file is converted from the time-decibel domain to the time-frequency domain. Then, the existing Mel filter bank is applied. This bank is a series of triangular filters spaced to best mimic human processing, with lower frequencies that have a higher resolution being represented more linearly, while higher frequencies with a lower resolution are represented more logarithmically. The resulting matrix is then normalized to reduce the computational cost in subsequent processing.

To further increase the robustness of our model, we also applied data augmentation techniques to the spectrograms. We randomly selected either a frequency mask or a time mask (Figure 3). Then, a randomly selected point in the spectrogram would have its value set to 0 for a duration that was also randomly selected between 0 and 20 milliseconds. For a frequency mask, a particular range of frequencies in the spectrogram is set to zero for the duration of the spectrogram. For the time mask, all the frequences for a particular time range is set to zero. In total, from each audio file (original and augmented), we generated one non-augmented spectrogram and three augmented ones^12^. Thus, we had a total of 14,848 spectrograms, which corresponds to 3,712 labeled groups, since there are 4 recordings per group.

## ALGORITHM DEVELOPMENT

### Vision Transformer

A Vision Transformer (ViT) is a deep-learning method that is effective in image recognition^13^ (Figure 4). A notable feature of ViTs is their patch-like approach to image classification. First, the input image is divided into patches of 16 × 16 pixels. Each patch is subsequently flattened into a 1-dimensional vector and adds a unique identifier to preserve the spatial orientation of the patch (called positional encoding). The patches with their respective positional encoding are passed into the transformer that utilizes a self-attention mechanism to assess and learn how the patches are related. During this process, a specialized token, called the [CLS] token, is added to the beginning of patch embedding sequences to gather information from all patches. Once transformer processing is complete, the [CLS] token will contain all critical information from the image. The [CLS] token is then passed to a dense neural network layer of the Transform Encoder, which will calculate a score for each preexisting category. These scores are termed logits and are still considered raw, unnormalized values. Conventionally, logits would undergo further processing to allow for classification of the image; however, we implemented ViT as an additional measure for image pre- processing, thus this step is bypassed.

**Figure 4.**
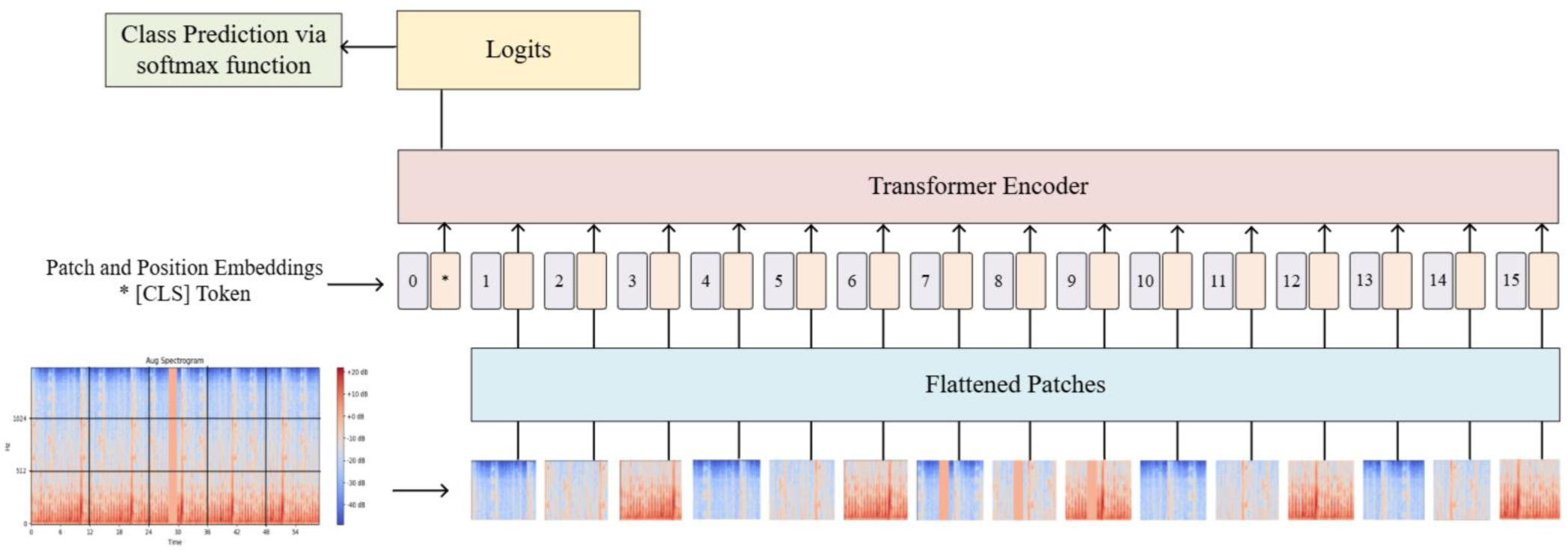
A visualization of the Vision Transformer methodology, beginning with patching and subsequent linear projection. Then, positional encoding is conducted, and the [CLS] token is added before inputting to the transformer. The transformer outputs a vector of logits that will be used for class prediction.

### MiniROCKET

The returned logits are then stored in a 2-dimensional structure similar to a table, called a DataFrame. This DataFrame contains two columns, one for the file name and one for the logit values. There is one row for each audio file. This data is then stored and reshaped into a 3- dimensional NumPy array called the X array. This array has the dimensions (groups, audio files, logits); thus its shape is (3712, 4, 1000). The ground truth diagnosis for each patient, either ‘murmur present’ or ‘murmur absent,’ is stored in a corresponding Y array, such that the diagnosis and data will be at the same index in their respective arrays.

The indices of the arrays are then randomly shuffled while preserving the X array and Y array compatibility. The data is then split into a training and testing dataset, with 80% of samples selected for the training set while the remaining 20% become the testing set. The training and testing sets contained both original and augmented data.

MiniROCKET makes use of a 1 × 9 matrix, called a kernel^6^. Each pixel in the kernel is assigned a weight of either -1 or +2 such that the sum of the kernel weights is equal to zero^6^. Each kernel is then passed over the data to calculate the dot product and generate a new signal of similar length to the original data (Figure 5). In a multivariate model, such as ours, each kernel is passed over each channel, corresponding to each heart valve, individually. Subsequently, the data is concatenated^6^.

**Figure 5.**
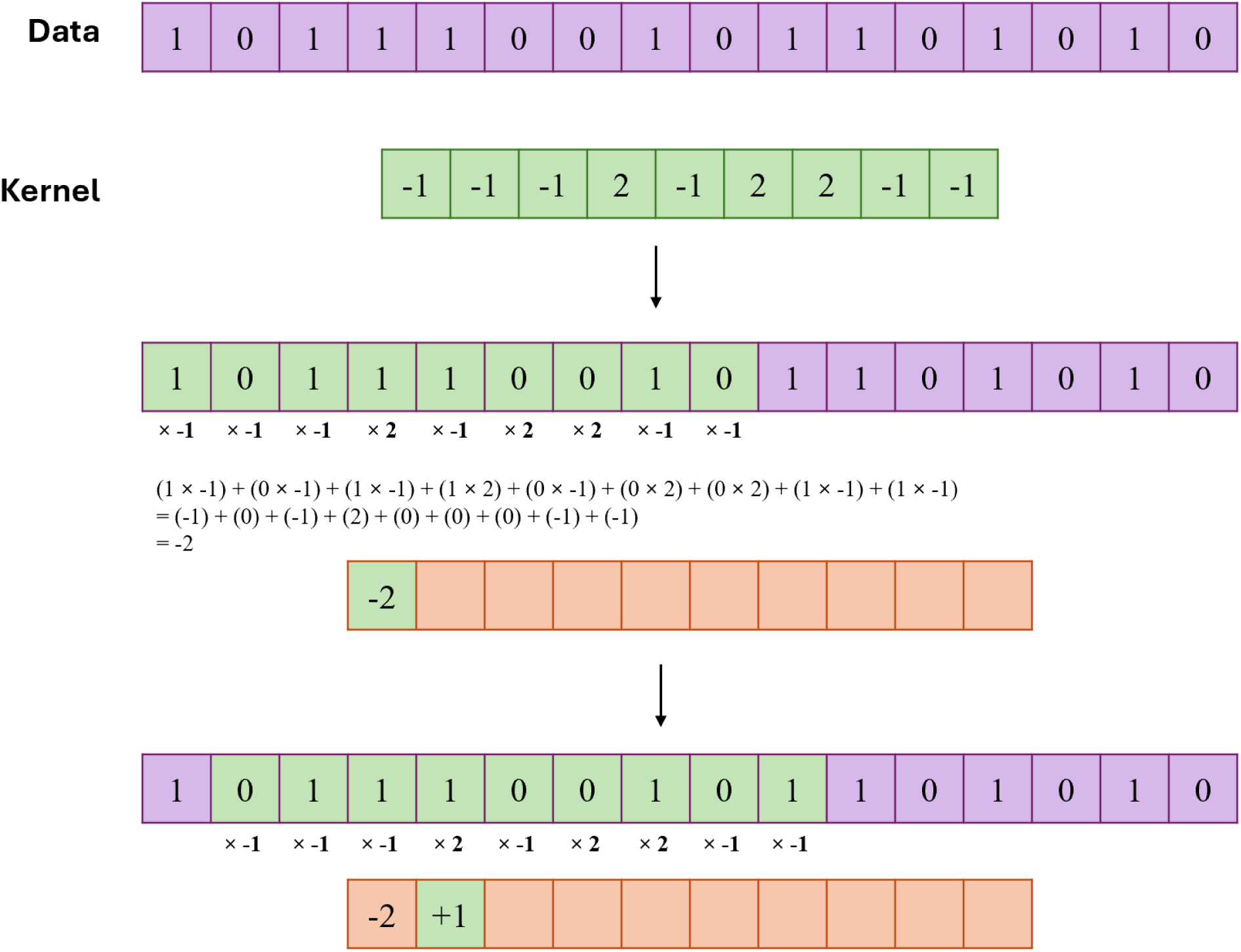
A visualization of dot product calculation through kernel convolution with a kernel size of 3x3 and a dilation rate of 1. The dot product is calculated as the sum of the products of each data pixel and the value of the corresponding kernel pixel. The kernel passes over the data to read it all in this fashion.

This process is completed with 10,000 distinct kernels to generate a feature map. This feature map then serves as the input for a Ridge Classifier, a specialized linear regression model that has been enhanced to prevent overfitting and increase generalizability during the classification process^14^. However, rather than utilizing the traditional ‘0’ threshold for class prediction, to improve confidence in the diagnosis, we propose a custom threshold of |0.1|. If the score calculated by the Ridge Classifier falls within this range, a prompt for “further review necessary” is returned, rather than definitively assigning a diagnostic class.

## STATISTICAL ANALYSIS

Once training and testing were completed, we assessed three aspects of our methodology:

1. The effect of varying levels of data augmentation
2. The overall accuracy of our MiniROCKET methodology
3. The speed of our model

This assessment was conducted by using numerous metrics to draw comparisons between our model and pre-existing models. Notably, we calculate Sensitivity, Specificity, Positive Predictive Value, Negative Predictive Value, F-Score, Accuracy and Weighted Accuracy. We also assessed the evaluation time of our model.

In our assessment, “True Positive” is synonymous with “Truly Present” and “True Negative” is synonymous with “Truly Absent”.

### F-Score

The F-Score provides an overall assessment of the algorithm’s functioning for each class (“present” or “absent”) with respect to Positive Predictive Value and Sensitivity, and is represented by the following formula [28]

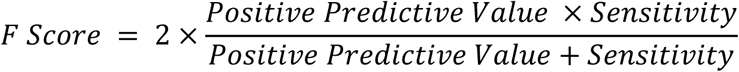

### Weighted Accuracy

Weighted Accuracy (WAcc) is a metric defined by the PhysioNet Challenge, which assigns a weight of 5 to True Positives to increase their significance in the accuracy calculation. It is represented by the following formula

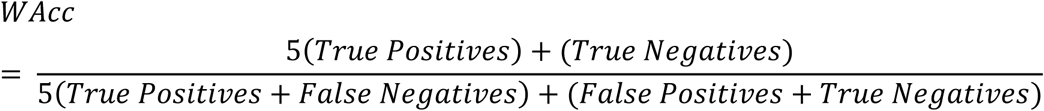

## RESULTS

Once training and testing were completed, the Confusion Matrices in Figure 6 were obtained. Each matrix was generated with increasing amounts of augmented training data. Subsequently, we conducted a quality metric analysis for each matrix to observe how these metrics change as sample size is increased. An Analysis of Variance (ANOVA) was used to assess the statistical significance of the observed improvements. Our statistical results are outlined in Table 1 and Table 2.

**Figure 6.**
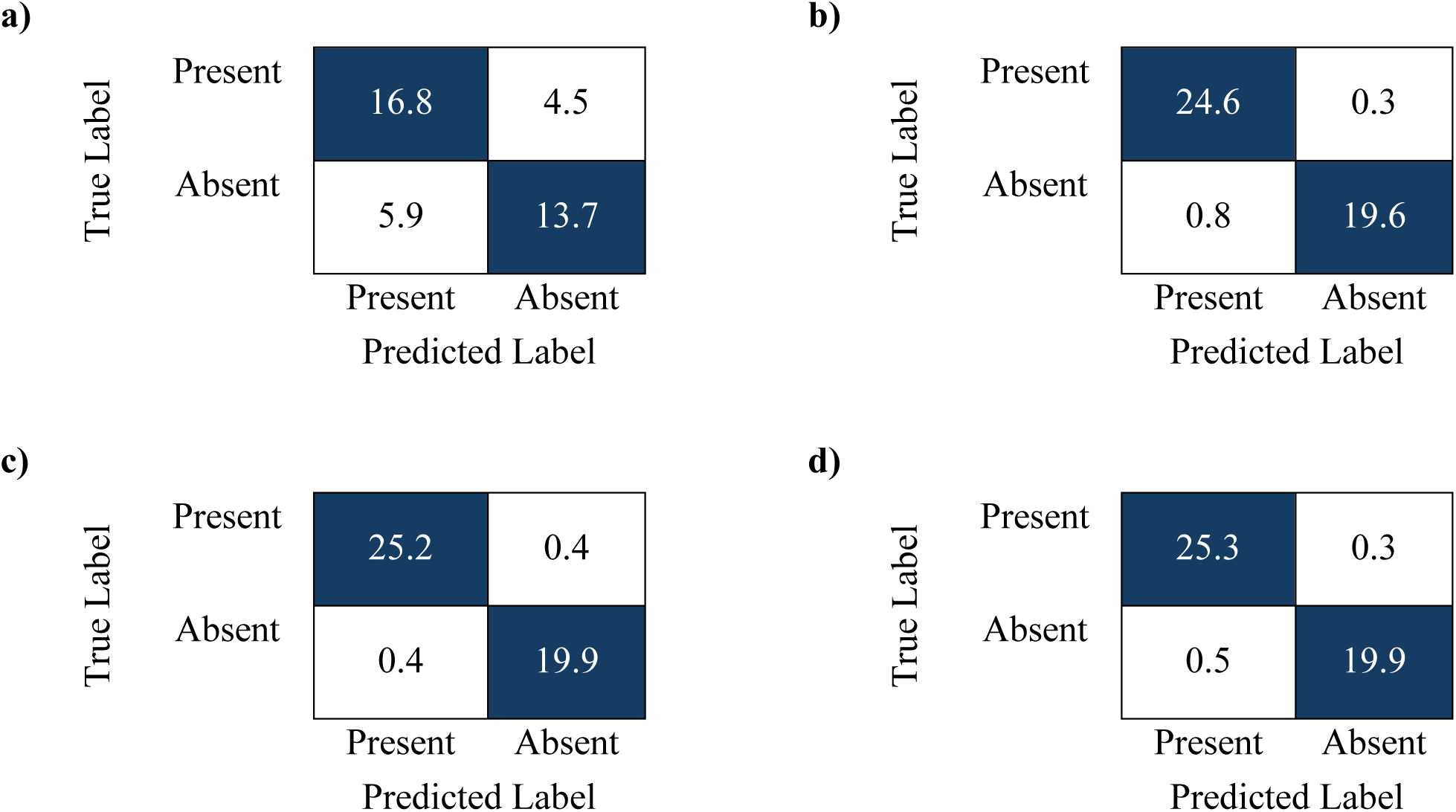
The confusion matrices that were generated during testing phases with various quantities of augmented data. **a)** The confusion matrix generated when no augmented data was used. **b)** The confusion matrix generated from original audio data and 1 augmented audio file, then 1 non-augmented spectrogram and 1 augmented spectrogram were generated from each original audio file for a total of 3,712 files. **c)** The confusion matrix generated when 2 augmented audio files and 2 augmented spectrograms were generated from each original audio file for a total of 8,352 files. **d)** The confusion matrix generated when 3 augmented audio files and 3 augmented spectrograms were generated from each original audio file for a total of 14,848 files.

**Table 1.**
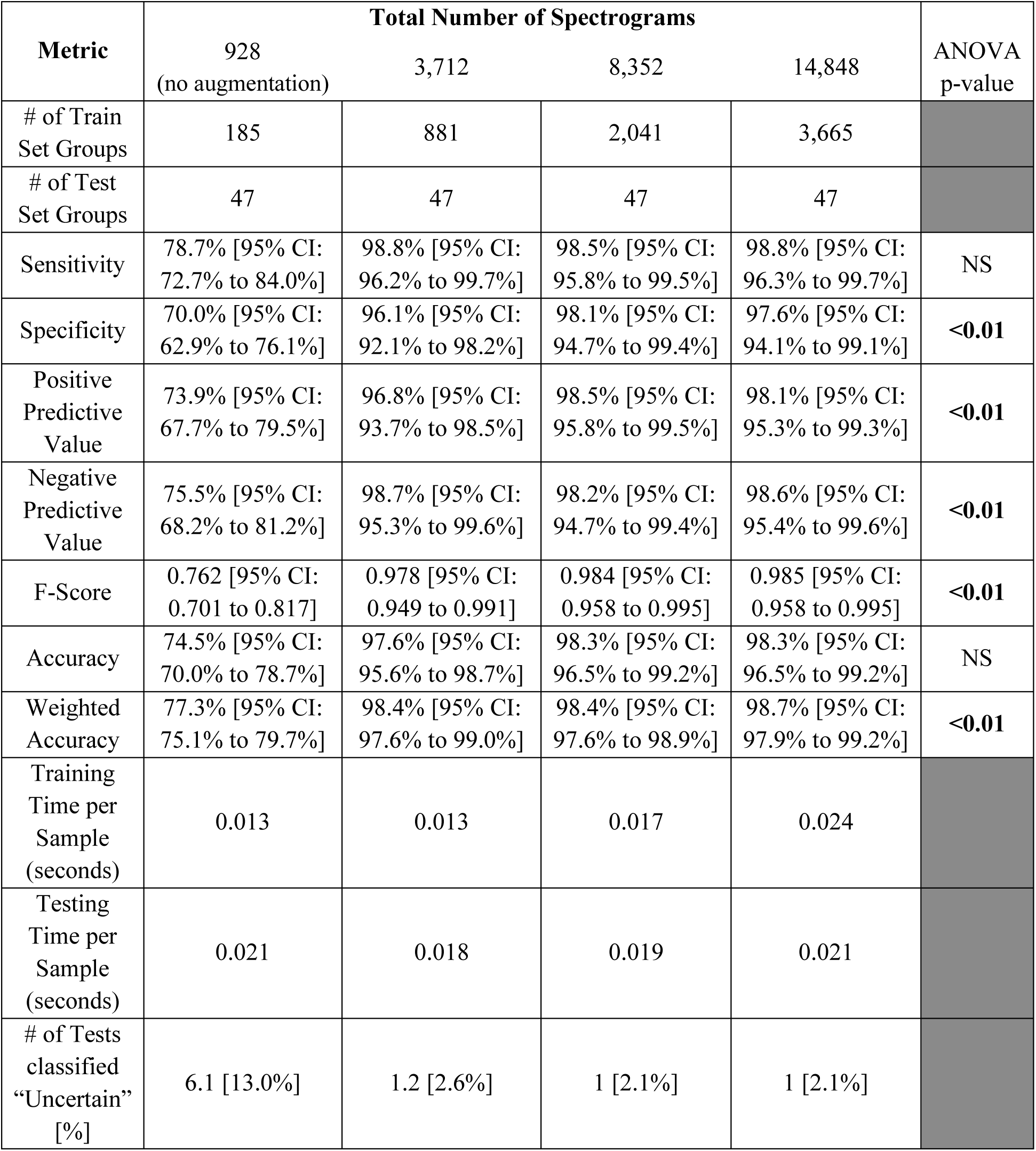
Comparing the performance of our model at various quantities of data augmentation.

**Table 2.**
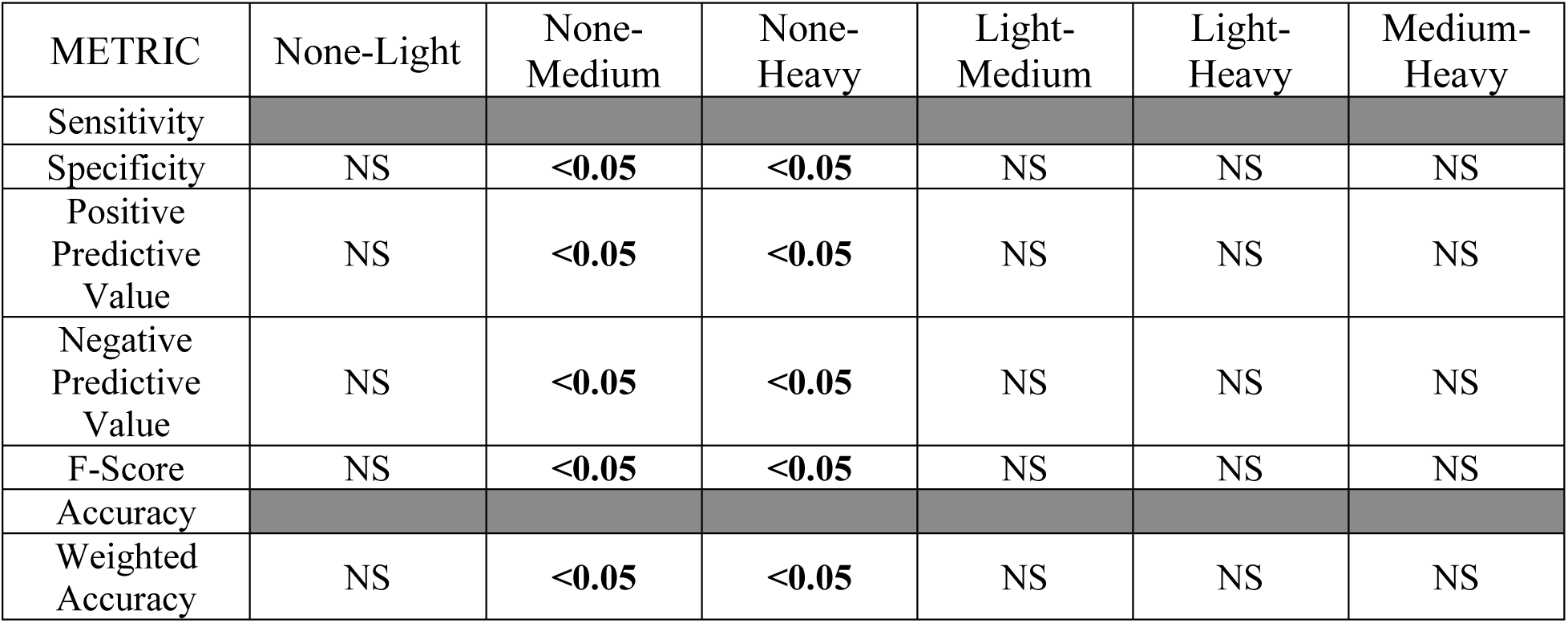
Post-Hoc Tukey Honestly Significant Difference analysis to determine significance of improvement in data augmentation.

| METRIC | None-Light | None-Medium | None-Heavy | Light-Medium | Light-Heavy | Medium-Heavy |
| --- | --- | --- | --- | --- | --- | --- |
| Sensitivity |  |  |  |  |  |  |
| Specificity | NS | <0.05 | <0.05 | NS | NS | NS |
| Positive Predictive Value | NS | <0.05 | <0.05 | NS | NS | NS |
| Negative Predictive Value | NS | <0.05 | <0.05 | NS | NS | NS |
| F-Score | NS | <0.05 | <0.05 | NS | NS | NS |
| Accuracy |  |  |  |  |  |  |
| Weighted Accuracy | NS | <0.05 | <0.05 | NS | NS | NS |

The results of the ANOVA yielded a statistically significant improvement in specificity, positive predictive value, negative predictive value, f-score, and weighted accuracy. Subsequently, Tukey’s Honestly Significant Difference was used to identify which specific groups have significant differences between them. The results, as observed in Table 2, demonstrate that all significant differences were observed between the no augmentation group and the groups with moderate and high levels of augmentation. Thus, increasing levels of augmentation contributes to overall enhancement of the classification model.

## COMPARISON WITH PRE-EXISTING MODELS

To assess the applicability of our model, we compare our metrics to pre-existing methodologies, notably the top 5 algorithms from the original 2022 PhysioNet Challenge, as well as the method introduced by Manshadi & Mihandoost in 2024^15^.

Manshadi & Mihandoost introduce a semi-supervised model for heart murmur detection in phonocardiogram data. Like us, they utilize data from the 2022 CirCor DigiScope Phonocardiogram Dataset. Their method begins by conducting a Stockwell transform to convert phonocardiogram data to 2-dimensional time-frequency maps^16^. Then, a deep convolutional neural network called AlexNet is used to extract deep features before subsequently passing them into a Support Vector Machine (SVM) Classifier. The metrics for their study are presented in Table 3, alongside the results of the top 5 algorithms from the original 2022 PhysioNet Challenge.

**Table 3.**
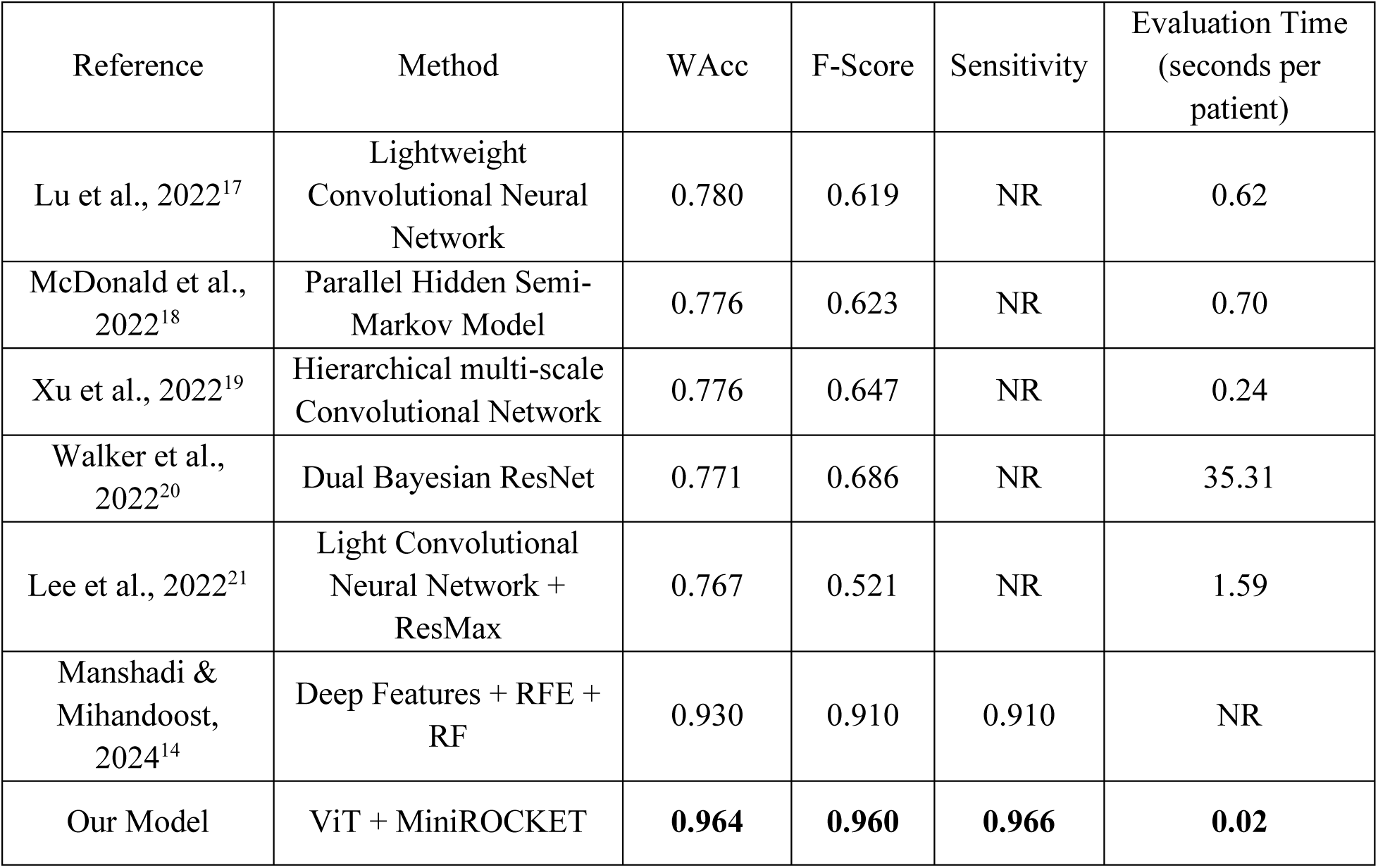
Comparing the performance of various models for heartbeat detection.

As depicted, our quality metric assessments demonstrate an enhanced murmur detection method when compared to all 2022 PhysioNet Challenge algorithms as well as the one proposed by Manshadi & Mihandoost^14^.

## EVALUATION TIME ASSESSMENT

A comparison with the PhysioNet Challenge algorithms highlighted our model’s enhanced efficiency, with our model spending roughly 0.02 seconds per patient during the evaluation phase, while the most efficient PhysioNet Challenge model, proposed by Xu et al., spent 0.24 seconds per patient^19^. Thus, our method is considerably more efficient.

With a Weighted Accuracy of 96.4%, Sensitivity of 96.6%, and F-Score of 0.960, as well as an evaluation time of 0.02 seconds per patient, our MiniROCKET implementation is superior compared to pre-existing methods.

## DISCUSSION

MiniROCKET provides several unique advantages. First, it has fewer trainable parameters compared to other machine learning methods, thus rendering it particularly effective for smaller datasets, enabling faster classification. Next, by using 10,000 convolution kernels, MiniROCKET strikes a balance between high accuracy and reduced computational costs, thus boasting improved classification efficiency^6^. Additionally, the utilization of random kernels prevents over-training and allows the model to focus on broad patterns in the data. Thus, it can effectively handle noise, making it especially robust for real-world applications^6^. Our results are further improved by leveraging data augmentation techniques to artificially increase the dataset size. The advantages of these techniques are evidenced by the results of our model. With a Weighted Accuracy of 96.4%, Sensitivity of 96.6%, and F-Score of 0.960, our method performs better on quality assessment metrics when compared to other existing methodologies. Furthermore, with an evaluation time of 0.02 seconds per patient, our MiniROCKET implementation completes heart murmur detection at a much faster rate than the top models from the PhysioNet Challenge^19^.

Faster algorithms provide notable advantages from both clinical and technical perspectives. Firstly, models with faster processing capabilities allow for quicker diagnostic feedback. This is particularly important in time-sensitive emergency situations. Additionally, early detection of underlying cardiovascular issues allows for earlier intervention and, thus, can promote improved long-term health outcomes. Furthermore, enhanced processing allows for more patients to be screened in a shorter period of time, thus allowing increased scalability and application in medical settings. From a technical perspective, shorter training and testing times make the model easier to adapt or refine when necessary. Updating the model is necessary to enhance its accuracy and robustness, as well as to ensure it is up-to-date and effective for its assigned task. Finally, reducing training and testing times also allows for a lower computational cost, making it less resource and energy-intensive. Increasing energy efficiency in this manner reduces the algorithm’s financial and environmental costs.

With recent medical and technological advancements, we have made strides in improving access to timely and accurate medical diagnosis. Yet, cardiovascular disease is still a critical cause for concern. Rapid and precise detection of murmurs in phonocardiogram data allows for sooner detection of underlying cardiovascular complications and thus allows early-stage treatment and improvements in long-term health outcomes. Furthermore, this method may allow for increased screening outside conventional healthcare settings, thus improving healthcare access.

Ultimately, this paper lays a foundation for future investigations regarding the implementation of MiniROCKET in heart murmur detection and classification. Since heart murmurs can be either benign or indicative of a more serious underlying condition, one critical next step will involve the development of a robust model capable of discriminating between these two categories. Additionally, this research opens the door for further explorations into the broader application of MiniROCKET in other medical domains. For example, future studies could investigate its efficacy in the analysis of electrocardiograms or electroencephalograms for anomaly detection, potentially revolutionizing medical diagnosis.

## ACKNOWLEDGEMENTS

We extend our sincere thanks to the peer reviewers for their time and thoughtful feedback, which greatly contributed to the improvement of this manuscript.

## DISCLOSURES

The authors declare that there are no actual or potential interests that relate to the research presented in this paper.

The authors attest that the primary author had full access to all data and takes responsibility for its integrity and data analysis.

## Non-Standard Abbreviations and Acronyms

MiniROCKET: Minimally Random Convolutional Kernel Transform
ViT: Vision Transformer
STFT: Short Time Fourier Transform

